# Opponent surrounds explain diversity of contextual phenomena across visual modalities

**DOI:** 10.1101/070821

**Authors:** David A. Mély, Thomas Serre

**Affiliations:** Department of Cognitive, Linguistic & Psychological Sciences; Brown Institute for Brain Science; Brown University, Providence, RI 02912, USA

**Keywords:** classical receptive field, extra-classical receptive field, visual cortex, illusion, induction, assimilation, contrast, tilt effect, enhanced perceptual shift

## Abstract

Context is known to affect how a stimulus is perceived. A variety of illusions have been attributed to contextual processing — from orientation tilt effects to chromatic induction phenomena, but their neural underpinnings remain poorly understood. Here, we present a recurrent network model of classical and extra-classical receptive fields that is constrained by the anatomy and physiology of the visual cortex. A key feature of the model is the postulated existence of two spatially disjoint near-vs. far-extra-classical regions with complementary facilitatory and suppressive contributions to the classical receptive field. The model accounts for a variety of contextual illusions, reveals commonalities between seemingly disparate phenomena, and helps organize them into a novel taxonomy. It explains how center-surround interactions may shift from attraction to repulsion in tilt effects, and from contrast to assimilation in induction phenomena. The model further explains enhanced perceptual shifts generated by a class of patterned background stimuli that activate the two opponent extra-classical regions cooperatively. Overall, the ability of the model to account for the variety and complexity of contextual illusions provides computational evidence for a novel canonical circuit that is shared across visual modalities.

---

Spatial context has been known to affect perception since at least Aristotle (Eagleman, 2001). The past several decades of work in visual psychophysics have revealed a plethora of seemingly disparate contextual phenomena whereby subtle differences in experimental conditions yield a wide variety of effects (Figure 1). In the classical tilt illusion (O’Toole and Wenderoth, 1977; Goddard et al., 2008), the perceived orientation of a center stimulus tilts either towards or away from that of a surround stimulus, depending on their relative orientations. Many variants have been tested with a variety of stimulus parameters including spatial frequency, color, luminance, contrast differences between center and surround stimuli as well as their spatial and temporal separation (see Clifford, 2014; for a review).

**Figure 1.**
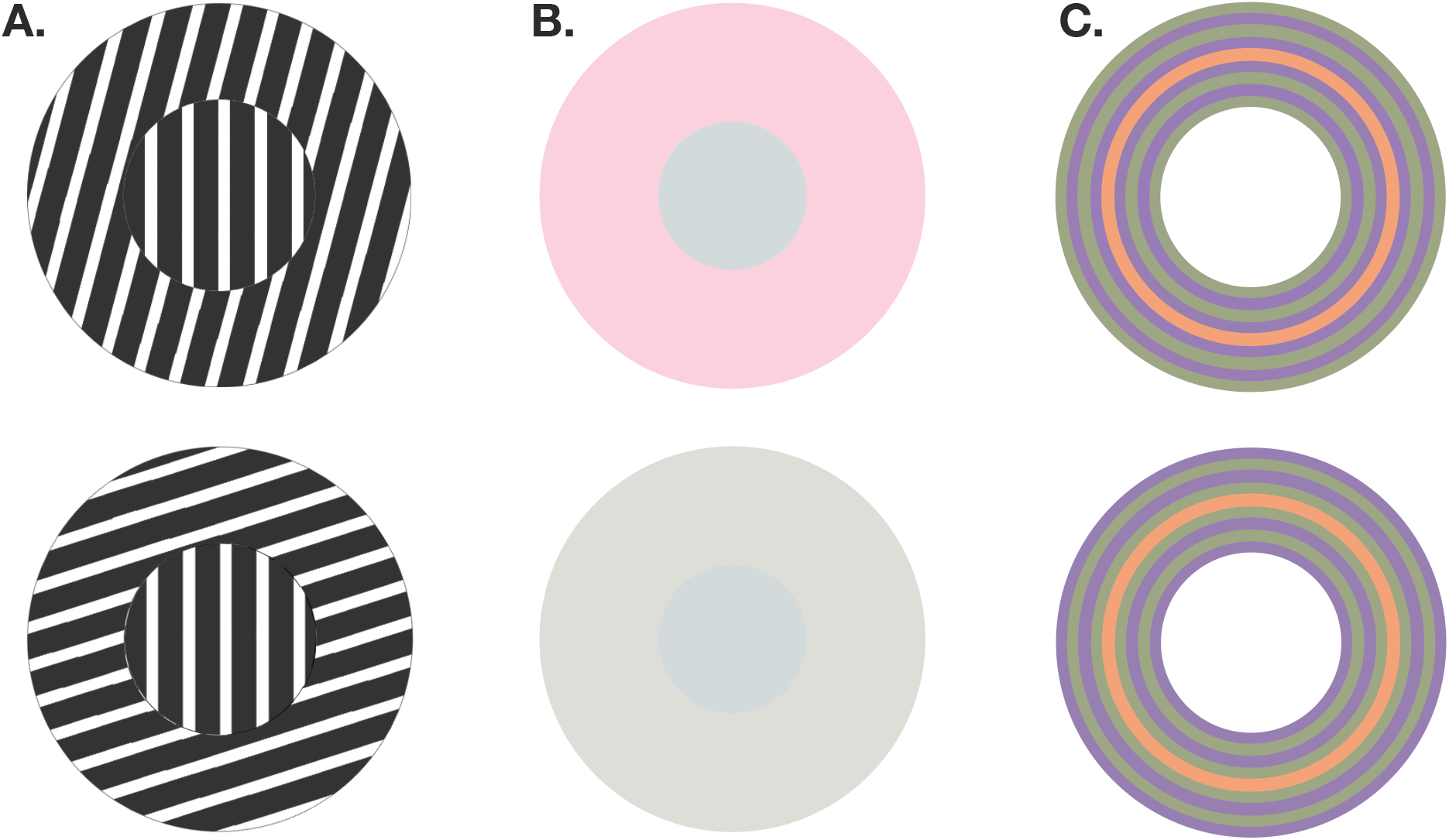
Representative contextual phenomena explained by the model. **A. Orientation tilt:** The perceived center (or test) orientation appears tilted from its true physical orientation, away from the surround (or contextual) orientation when center and surround stimuli are similar (top) and towards the surround orientation when they are dissimilar (bottom). **B. Color induction:** A central gray stimulus appears greener when embedded in a pink surround (top) compared to a neutral gray surround (bottom). **C. Enhanced color shifts:** The test stimulus is a central, orange ring, embedded in a surround stimulus composed of alternating purple and lime rings. The test ring looks vividly more pink when the adjacent color is purple, followed by lime (top), and looks more yellow when lime is the adjacent color, followed by purple (bottom).

Similar effects have been reported in the motion domain – for both direction and speed (Marshak and Sekuler, 1979; Murakami and Shimojo, 1993; 1996; Kim and Wilson, 1997). In color induction, both the spatial frequency and phase of the surround controls the direction of the perceived shift in hue of a center stimulus relative to that of the surround (Smith et al., 2001; Monnier and Shevell, 2003; Shevell and Monnier, 2005). In the disparity domain, a center stimulus appears closer or further away from an observer, depending on the relative depth and spacing between center and surround stimuli (Westheimer, 1986; Westheimer and Levi, 1987). While much is known about the psychological basis of these phenomena, our understanding of the underlying neural mechanisms remains, at best, fragmentary.

A widely held assumption is that such contextual phenomena are mediated in the cortex by extra-classical receptive field (eCRF) mechanisms (reviewed in Seriès et al., 2004; Angelucci and Shushruth, 2013): The presentation of a stimulus in the eCRF alone does not typically elicit any response from the cell but modulates its response to a stimulus presented in the classical receptive field (CRF). Such center-surround interactions have been reported across visual modalities including orientation and spatial frequency (DeAngelis et al., 1994), motion (Li et al., 1999; Jones et al., 2001), color (Schein and Desimone, 1990; Wachtler et al., 2003) and disparity (Bradley and Andersen, 1998).

Although several eCRF models have been developed to describe specific phenomena (reviewed in Seriès et al., 2004; Angelucci and Shushruth, 2013; see also Discussion), a unifying theory, which would integrate disparate aspects of contextual integration and, ultimately, link primate neurophysiology to human behavior, is still lacking. We have thus developed a large-scale recurrent network model of classical and extra-classical receptive fields that distinguishes itself from previous work – allowing us to simulate realistic cortical responses to a variety of full-field, real-world, stimuli (including center-surround stimuli) defined across visual modalities (we model orientation, color, motion, and binocular disparity). The model is constrained by anatomical data and shown in our experiments to be consistent with V1 neurophysiology. A key feature of the model is the postulated existence of two spatially-disjoint near vs. far extra-classical eCRF regions with complementary asymmetric contributions (facilitatory vs. suppressive) to the CRF response. Using an ideal neural observer, we show that the model is consistent with human behavioral responses for a variety of contextual phenomena – revealing commonalities between seemingly disparate phenomena and helping to establish a novel taxonomy of contextual illusions.

## Results

The visual cortex is modeled as a dense, regular topographic grid of cortical (hyper)columns which tile the visual field (Figure 2A). Each hypercolumn contains a complete set of units with coinciding CRFs. Their tuning curves are idealized (see Materials and Methods), and centered at regular intervals (e.g., between 0° and 180° for orientation-tuned units). For simplicity, we do not take into account cortical magnification and assume a fixed sampling of the visual field at all eccentricities. The model takes into account connections both within and across hypercolumns in order to explain several CRF and eCRF properties. The resulting circuit motif is replicated around every hypercolumn.

**Figure 2.**
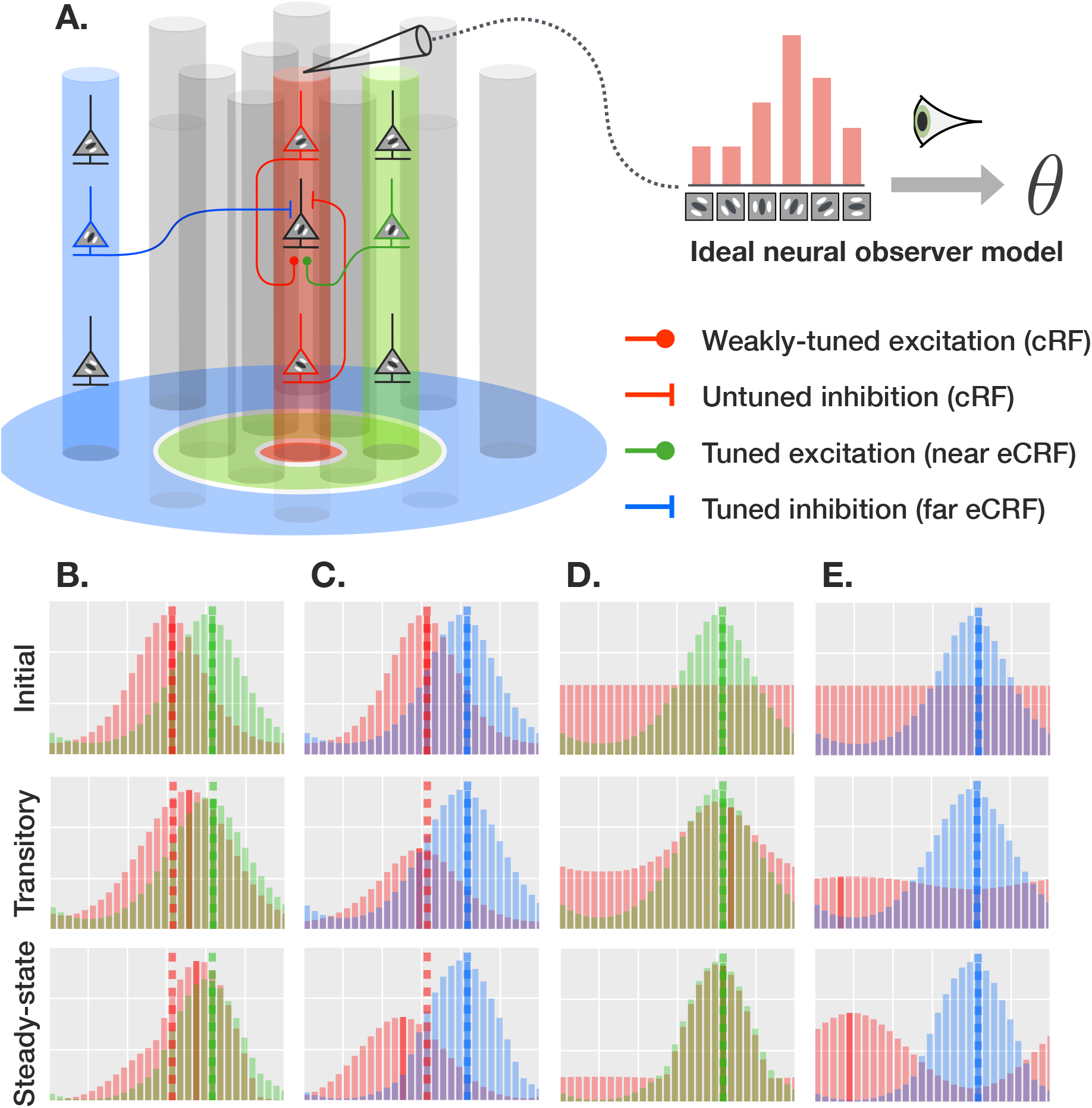
Recurrent network model of center-surround interactions. **A. Connectivity:** The model implements excitatory and inhibitory connections both short-range (within hypercolumns) and long-range (between hypercolumns). The regions shown in red, green and blue correspond to the CRF, near eCRF and far eCRF, respectively (defined for the reference column in red). Model inhibitory connections are such that the net inhibition onto a target unit is a function of not just the pre-synaptic activity, but also the post-synaptic activity (see text for details). Color conventions for the CRF and the near and far eCRFs are used consistently throughout the paper. **B–E. Representative model dynamics:** Example population responses (32 direction-tuned model units) following the presentation of a contextual stimulus corresponding to the initial, transitory and steady state (rows). Population responses correspond to locations in the CRF, near eCRF and far eCRF. Highlighted bars represent directions decoded from the corresponding populations (undefined for flat responses); dashed lines represent initial decoded values at stimulus onset. Each column corresponds to a representative transformation undergone by the center population under the proposed taxonomy of contextual phenomena derived from the model: **B.** attractive shift, **C.** repulsive shift, **D.** bump, **E.** notch. (see also Discussion and Figure 8). Abscissas span the range [−180°, 180°] and ordinates are normalized independently for readability.

### Intra-columnar recurrent circuits

Recurrent connections (Figure 2A, red connections) within a column (i.e., originating from within the CRF) include both local excitatory and inhibitory connections. Inhibitory CRF contributions constitute one of the key mechanisms in an influential model of gain control (divisive normalization, reviewed in Carandini and Heeger, 2012). This model accounts for cross-orientation normalization phenomena (when a grating stimulus is masked by another one at any orientation, see Heeger, 1993; Carandini and Heeger, 1994) and was later extended to capture neural population responses, in order to account for the competitive interactions within a single hypercolumn (Busse et al., 2009; Sit et al., 2009). A recent optogenetic study demonstrated that the underlying circuits are recurrent rather than feedforward (Nassi et al., 2015).

Because this form of suppression does not seem to depend on the orientation of the afferent and target cells, it is often called “untuned” inhibition. In our model, we speculate that such untuned inhibitory recurrent connections within hypercolumns exist for all other visual domains (including color, motion as well as binocular disparity).

In addition to short-range inhibitory connections within hypercolumns, the model also incorporates short-range excitatory connections. In the cortex, such excitatory connections may drive neurons up to ten times more strongly than their feed-forward inputs (Douglas et al., 1995; Stepanyants et al., 2008). As suggested by Shushruth et al. (2012), we have found that recurrent excitation within the CRF is essential to account for some of the more complex aspects of surround suppression (see Supplementary Experiments, Figures S5 and S6), by placing the column in a regime dominated by recurrent as opposed to feed-forward inputs. Experimental data on the selectivity of these recurrent excitatory connections are scarce. Here, we assume that the corresponding local excitatory connections within a hypercolumn are only weakly tuned, as perfectly untuned excitation would effectively “flatten out” population response curves.

### Inter-columnar recurrent circuits

Expanding the optimal stimulus of a cortical neuron immediately beyond its CRF may facilitate its response (Bringuier et al., 1999; Sengpiel et al., 1997; Sceniak et al., 1999; Angelucci et al., 2002a;b; Briggs and Usrey, 2011). The corresponding eCRF region immediately surrounding the CRF integrates sub-threshold responses to surround stimuli that do not elicit any action potential when presented alone (Bringuier et al., 1999). It is thus a region considered to be distinct from the CRF.

A potential neural substrate for the near eCRF includes the short-range, tuned excitatory networks (Lee et al., 2016) which span a spatial extent consistent with that of the eCRF facilitation (Angelucci et al., 2002a;b) and amplify co-occurring local inputs at similar orientations (Sengpiel, 1997; Sceniak et al., 1999; Angelucci et al., 2002a;b; Briggs and Usrey, 2011). Because it is located immediately beyond the CRF, we deem this region the *excitatory near eCRF* (or near surround; green annulus and connections in Figure 2A). In the model, we assume that all excitatory connections from other hypercolumns centered in the near surround are tuned, irrespective of the visual modality (i.e., the stimulus with the preferred orientation, or direction of motion, etc. in the CRF is also most effective in the near eCRF).

Expanding the optimal stimulus beyond the near eCRF results in neural suppression (first reported by Hubel and Wiesel, 1968; as hypercomplex tuning). Critically, the presentation of the suppressing stimulus in the eCRF alone does not elicit any activity from the recorded cell (see Angelucci and Shushruth, 2013; for review). The tuned nature of these suppressive mechanisms is well documented across visual modalities: from orientation (Hubel and Wiesel, 1968; DeAngelis et al., 1994; Weliky et al., 1995; Petrov et al., 2005; Ozeki et al., 2009) to color (Schein and Desimone, 1990; Wachtler et al., 2003), spatial frequency (DeAngelis et al., 1994), temporal frequency (Li et al., 1999; Jones et al., 2001), motion direction and speed (Allman et al., 1985) as well as binocular disparity (Bradley and Andersen, 1998).

Thus, we also define an *inhibitory far eCRF* (or far surround; blue annulus and connections in Figure 2A), located immediately beyond the near eCRF. In our model, a hypercolumn receives tuned inhibition from hypercolumns centered in its far surround.

To summarize, contributions from the eCRF as a whole arise from spatially segregated regions with opposite polarities. We do not assume any gap between the CRF and the near eCRF, nor between the near eCRF and the far eCRF. The first key assumption of the model is that, unlike local recurrent interactions within a hypercolumn, interactions across hypercolumns are “tuned” as only units that share the same preferred stimulus are directly connected.

The second key assumption of the model is an asymmetry between excitation and inhibition: In the model, excitation only depends on pre-synaptic activity and is purely additive. Inhibition, on the other hand, from either the CRF or eCRF, depends on both pre-and post-synaptic activity, and ultimately results in a combination of subtractive and divisive effects. Similar forms of inhibition have been used in previous recurrent network models to achieve divisive normalization. In practice, this means that, given a fixed amount of pre-synaptic inhibition, weakly active units receive less effective inhibition than more active ones. In contrast, any given amount of pre-synaptic excitation results in the same amount of effective post-synaptic excitation.

### Neural field model

Upon the presentation of a stimulus, recurrent interactions between units yield complex model dynamics. In particular, population responses at any location are modulated first by their immediate (near and far) eCRFs, then by responses across the visual field as transient activity propagates through the network, until all unit responses settle into a steady-state. Such short and long-range interactions are modeled using coupled differential equations and the steady-state solution of the resulting neural field model is computed using numerical integration methods (see Materials and Methods).

Next, we describe experiments conducted *in silico* to compare model responses to published psychophysics data. Psychophysics studies typically record perceptual judgements related to a center stimulus under varying surround conditions. To approximate these judgements, we use an ideal neural observer which maps center population responses to a sensory value. Note that in most cases, several columns may be located within the center stimulus; while any of these columns would be suitable for readout by the ideal observer, we selected the center-most column for simplicity (unless specified otherwise). Surround modulation thus gets translated into measurable perceptual changes in the center that can then be compared to human behavioral data.

We have organized these experiments into three broad categories, each reflecting a key computational mechanism and highlighting commonalities across visual modalities. As we will show, these experiments allow a clear picture to emerge: The diversity of observed contextual phenomena may result from a balance between two opposing “forces” that arise from complementary near-far eCRF mechanisms with opposing polarities. Figure 2 shows examples of CRF and eCRF population responses recorded from the model together with representative transformations they undergo as a result of these two forces (see Discussion for more details).

All model parameters (Table S1) governing the dynamics and relative strengths of the interactions between the CRF and the eCRF sub-regions were initially adjusted for the model to reproduce a host of V1 neurophysiology data (see Supplementary Experiments) including a comparison with data from Busse et al. (2009) (see Figures S3 and S4) and Trott and Born (2015) (see Figures S5 and S6). They were held fixed for all subsequent comparisons with psychophysics data. After scaling the stimulus, the only model parameter that was optimized for individual modalities was the tuning bandwidth of individual model units.

### Competitive activation of the near vs. far surrounds

Our comparison between experimental and model data starts with a set of three experiments that span the orientation, motion and color domains. All experiments involve simple center-surround stimuli, in which the surround stimulus is expected to jointly activate the near and far eCRFs. Thus, these experiments should reveal a fundamental aspect of the model: the outcome of a competition between the near facilitatory and the far suppressive eCRFs when they are simultaneously activated by a surround stimulus.

The orientation tilt occurs when the perceived orientation of a center stimulus is biased either towards or away from the orientation of a surround stimulus, also called the inducing stimulus or inducer (O’Toole and Wenderoth, 1977; Goddard et al., 2008). Figure 3C shows representative psychophysics data (digitally extracted from Figure 4 in O’Toole and Wenderoth, 1977; only data averaged across subjects are available from that study). These data are characterized by two regimes: a *repulsive* regime (i.e., the perceived center orientation shifts away from the surround orientation, corresponding to positive ordinates) when the surround orientation is similar to that of the center and an *attractive* regime (i.e., the perceived center orientation shifts towards the surround orientation, corresponding to negative ordinates) when the surround orientation is different enough from that of the center.

**Figure 3.**
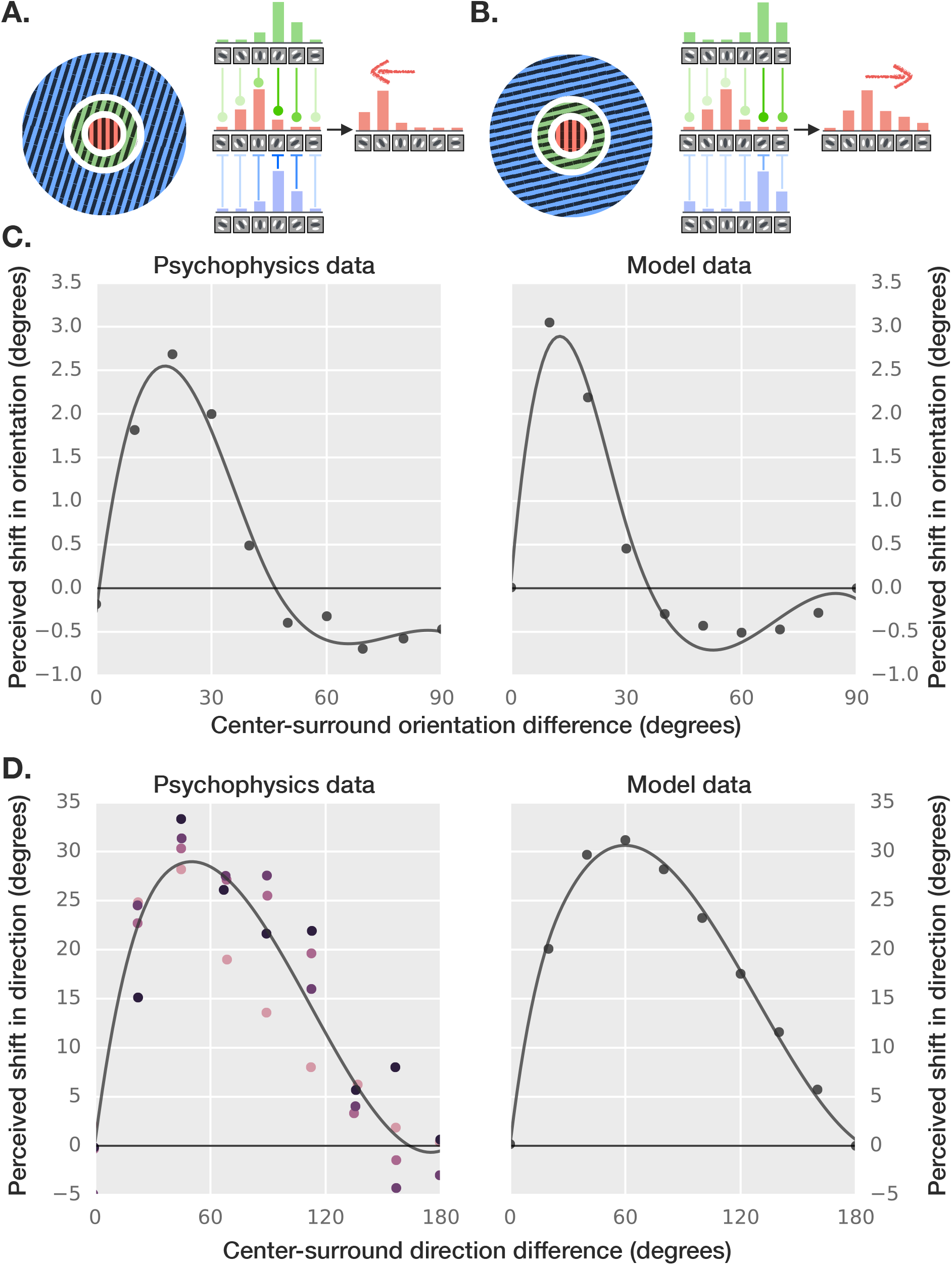
Tilt effects. Competitive activation of the near vs. far eCRFs explains the shift in tilt from one direction to the other. **A. Repulsion:** For similar center-surround orientations, tuned inhibition from the far eCRF outweighs excitation from the near eCRF, which yields a net repulsive force on the center population responses (away from that the surround orientation). **B. Attraction:** For dissimilar center-surround orientations, tuned excitation from the near eCRF prevails, which yields a net attractive force on the center population responses (towards the surround orientation). Note that gaps between the CRF and the near and far eCRFs were added for improved readability only and are not present in the actual model. **C. Orientation tilt: Psychophysics vs. model data.** Psychophysics data were digitally extracted from Figure 4 in (O’Toole and Wenderoth, 1977) and fitted with splines. The model explains the characteristic shift from perceptual repulsion (positive ordinates) to attraction (negative ordinates). **D. Motion tilt: Psychophysics vs. model data.** Psychophysics data were digitally extracted from Figure 3 (“periphery” condition) in (Kim and Wilson, 1997) and fitted with splines. Different colors correspond to different subjects. Both psychophysics and model data exhibit a similar dependency on the direction difference between center and surround, as well as a lack of an attractive regime.

**Figure 4.**
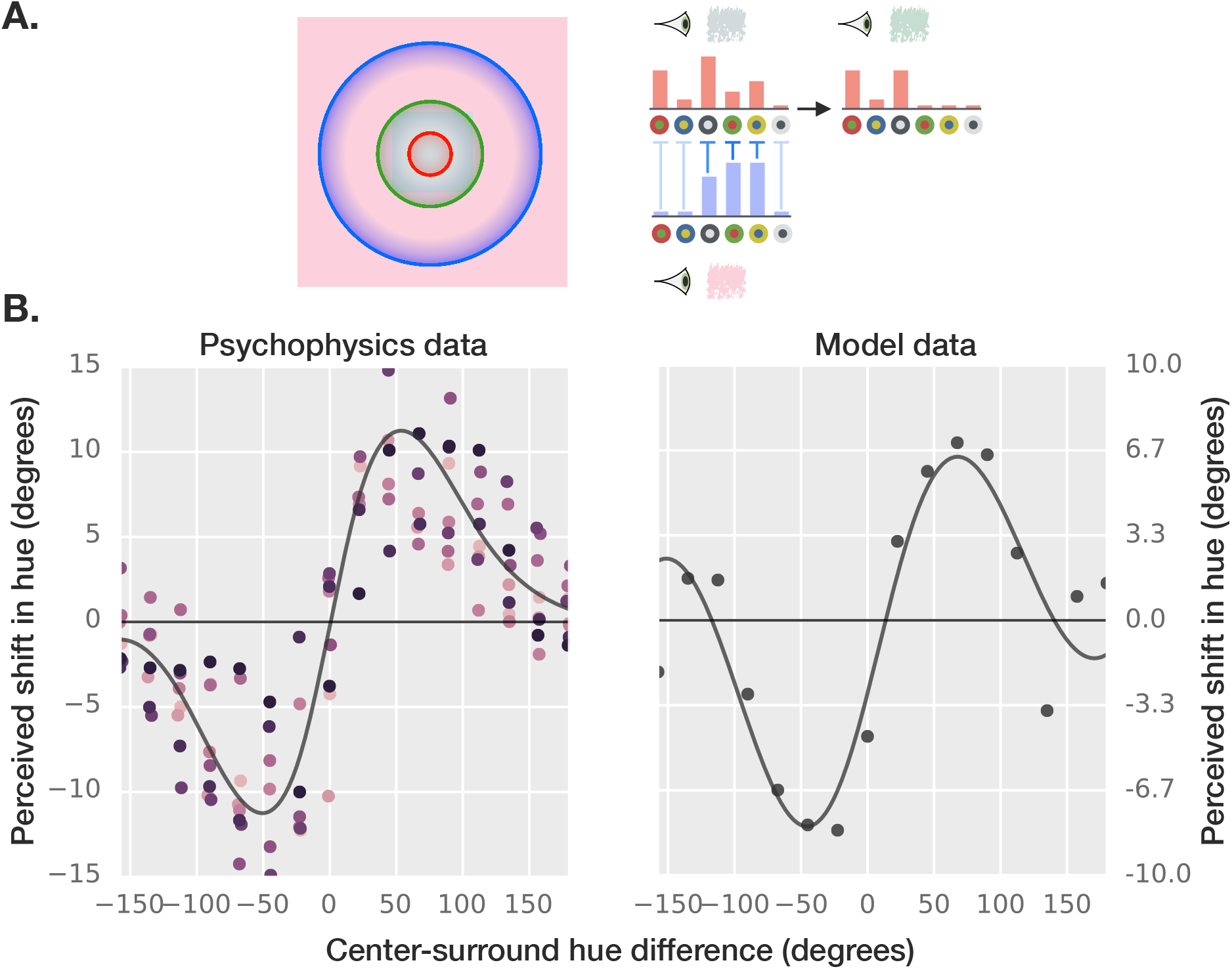
Color induction (or hue tilt effect). This experiment generalizes the tilt effect to opponent population codes. **A. Repulsion:** As with the classical tilt effect, the key model mechanism behind perceptual repulsion is the tuned inhibition from the far eCRF. In this example, the pink surround suppresses “red” center neurons, therefore reducing the “redness” of the gray center patch yielding a shift in the perceived center hue towards green. The same explanation also applies to chromatic center stimuli. Colored patches shown next to the eyes correspond to the color decoded under the ideal observer. **B. Psychophysics vs. model data:** Psychophysics data were digitally extracted from Figure 2 in (Klauke and Wachtler, 2015) and fitted with splines (averaged across eight surround hues). Both model and behavioral data exhibit a characteristic two-lobed shape peaking around ±50°.

The model successfully reproduces this balance between attraction and repulsion (Figure 3C; a similarly good fit was also obtained using broadband oriented textures as done in Goddard et al., 2008; data not shown). The key mechanism which enables the emergence of these two regimes is the postulated asymmetry between facilitatory and suppressive interactions originating from the near and far eCRFs, respectively. The net inhibition in the model, unlike excitation which is only dependent on pre-synaptic activity, increases monotonically with the level of post-synaptic activity of a target unit.

As a result, when neural population responses in the CRF and eCRF overlap significantly (as when center and surround orientations are similar), inhibition predominates and center population responses get comparatively more suppressed at orientations close to that of the surround. The center of mass of center population response curves shifts away from the surround orientation, biasing the neural decoding accordingly (Figure 3A). The surround thus acts as a repellent in this regime. In contrast, when neural population responses in the CRF and eCRF are far more offset (as when the surround orientation is near orthogonal to that of the center), excitation from the near eCRF predominates, and increases the activity of center units selective for the surround orientation. This results in a force that pushes the center population response towards the surround orientation. This, in turn, biases the decoding of the center orientation in the direction of the surround orientation (Figure 3B). The surround thus acts as an attractor in this regime.

Beyond the orientation domain, tilt effects have also been reported for the perception of motion direction. Figure 3D shows representative psychophysics data (digitally extracted from Figure 3 “periphery” condition in Kim and Wilson, 1997). Unlike in the orientation domain, however, perceptual shifts are always repulsive (the perceived motion direction of the center grating tilts away from that of the surround grating; both gratings have the same contrast and speed). This phenomenon can also be induced using coherently moving random dots (Marshak and Sekuler, 1979); the effect seems to peak for similar center-surround differences in motion direction (between 40° and 60°) for either kind of stimuli.

We found both a qualitatively and quantitatively good fit between the model and psychophysics data as shown in Figure 3D. In the model, the disappearance of the attractive regime is accounted for by a broadening of the tuning curves (compared to orientation; see Supplementary Materials and Methods). Interestingly, this seems consistent with neurophysiology data from the primary visual cortex (Ringach et al., 2002; Albright et al., 1984).

In our next experiment, we show that the model is also able to account for tilt effects in the hue domain, more widely known as color induction. The model reproduces the known shifts in human judgment obtained when a center hue is surrounded by an isoluminant background of a different hue (digitally extracted from Figure 2 in Klauke and Wachtler, 2015; averaged across multiple combination of center-surround hues sampled uniformly and independently as done in the original experiment).

As for motion induction, only a repulsive tilt effect is observed with hue. The model’s ability to account for these data is evident from Figure 4B, which confirms the hypothesis by Klauke and Wachtler (2015) that color induction is in fact just another tilt effect (i.e., a “hue tilt effect”). Furthermore, the same mechanisms that are responsible for the tilt effect in the orientation and motion domains, namely the balance between facilitatory and suppressive forces originating from the eCRF, are also at play in color induction (Figure 4A). However, opponent color coding yields populations from the center and the surround with high overlap, which explains the absence of an attractive regime for this phenomenon.

### Exclusive activation of the near vs. far surrounds

Previous experiments involved stimuli that reflected the outcome of a competition between the near and far eCRF, which were activated jointly. Here, instead, we consider experiments that are based on surround stimuli that activate the near or the far eCRFs separately.

In classical depth induction experiments (Westheimer, 1986; Westheimer and Levi, 1987), human observers are presented binocularly with a center test stimulus (e.g., a thin bar) flanked by two surround stimuli (e.g., parallel thin bars or small squares). The disparities of the flanker stimuli are adjusted so that they appear in the same depth plane, either slightly in front of or behind the center stimulus. The planar separation between the center and flanker stimuli (i.e., their distance in the fronto-parallel plane) is varied systematically. Examples where the flankers appear behind the center stimulus for a shorter and a larger separation are shown in Figure 5A-B).

**Figure 5.**
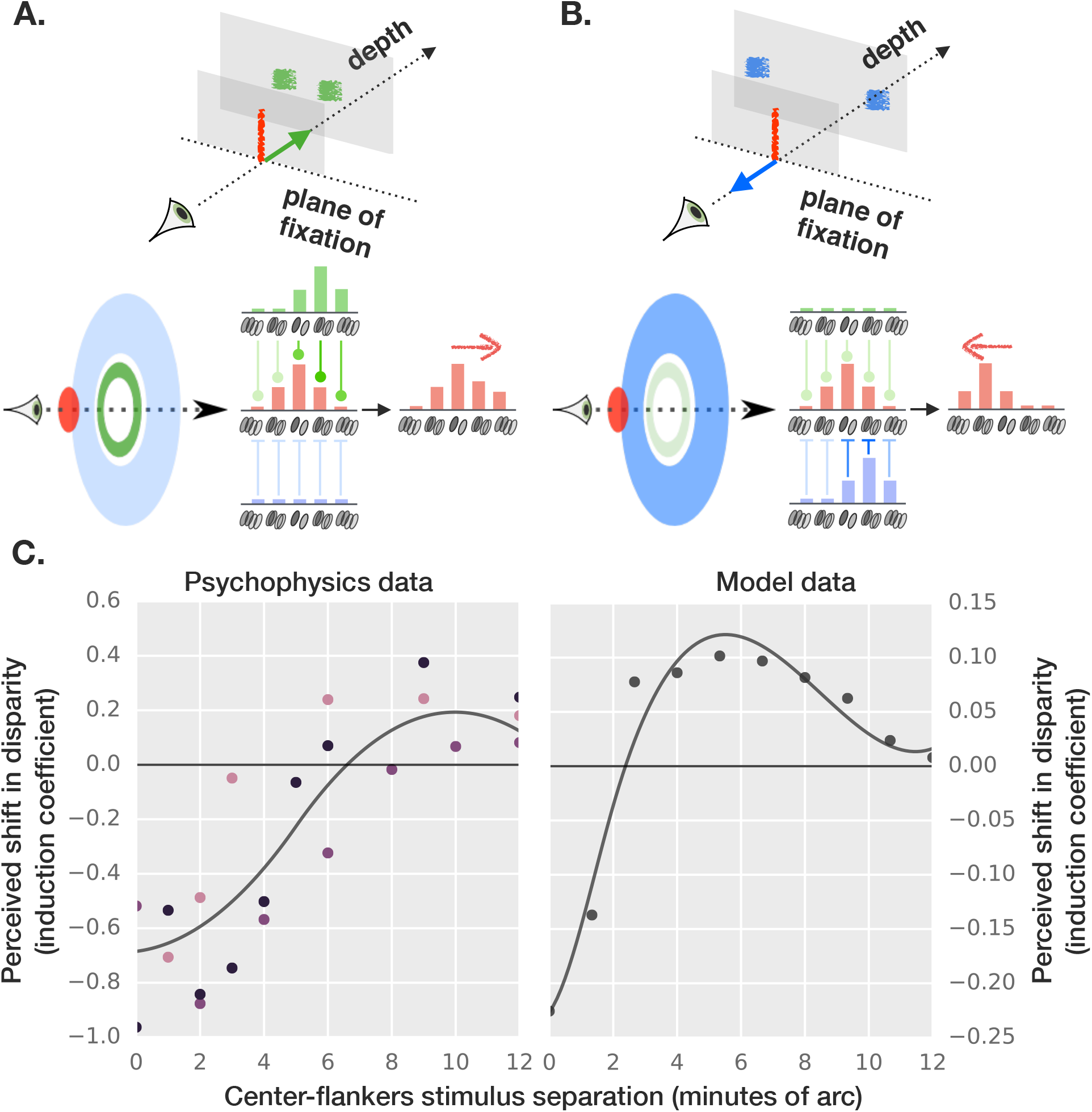
Depth induction. The exclusive activation of either the near or far eCRFs by flankers (as their separation vary) explains the existence of the shift from assimilation to contrast. The perceived depth of a binocular center stimulus (at zero disparity) is affected by binocular flankers located on either sides, presented at either crossed or uncrossed disparities. **A. Attraction:** For short separations, flankers activate the near eCRF, which yields a net attractive force on the center population responses (towards the surround disparity) corresponding to a negative perceived shift. **B. Repulsion:** For larger separations, flankers activate the far eCRF, which yields a net repulsive force on the center population responses (away from the surround disparity) corresponding to a positive perceived shift. **C. Psychophysics vs. model data:** Psychophysics data were digitally extracted from Figure 1 (upper panels) in (Westheimer and Levi, 1987) and fitted with splines. Both behavioral and model data capture the balance between stronger attraction towards the flankers at small separations, and weaker repulsion at larger separations. Note that the agreement between the model and human data is only qualitative as the perceived shifts in disparity are on different scales (the model underestimates the strength of the attractive regime in this illusion).

Results from the original study (data digitally extracted from Figure 1, upper panels in Westheimer and Levi, 1987) are shown in Figure 5C. When the flankers are close enough to the center stimulus, they seem to attract it in depth (corresponding to a negative shift in perceived disparity for very small flankers/center separations). That is, the center stimulus appears closer to (further away from) the observer when the flankers are in front of (behind) the center stimulus. Instead, when the flankers are moved far enough laterally, they start to repel the center stimulus (corresponding to a positive shift in perceived disparity for larger flankers/center separations).

The observed shifts in depth found in the model (Figure 5C) matches qualitatively with human psychophysics data: Flanker stimuli located close enough to the test stimulus activate solely the near eCRF, resulting in a purely facilitatory net eCRF influence. As with the aforementioned tilt effects, net facilitatory eCRF contributions yield attraction of the center towards the surround. Conversely, flankers that are far enough from the test stimulus activate solely the far eCRF. This results in a net suppressive eCRF influence, which translates into repulsion of the center away from the surround.

A classical stimulus used in motion direction induction (Murakami and Shimojo, 1993; 1996) includes a center which consists of random moving dots devoid of coherent motion overall. The addition of a surround consisting in random dots coherently moving in one direction elicits the perception of illusory coherent center motion – either in the same or in the opposite direction to that of the surround. In (Murakami and Shimojo, 1993; 1996), the scale of the center-surround stimulus was systematically varied while the center and surround region sizes were held in constant proportion.

Figure 6C shows psychophysical data (digitally extracted from Figure 6 and 7 in Murakami and Shimojo, 1996). For small stimulus sizes, the induced center movement is in the same direction as that of the surround. This corresponds to an attractive regime measured as a negative shift in the point of subjective equality (PSE). For larger sizes, the center induced movement reverses direction. This corresponds to a repulsive regime measured as a positive shift in PSE.

**Figure 6.**
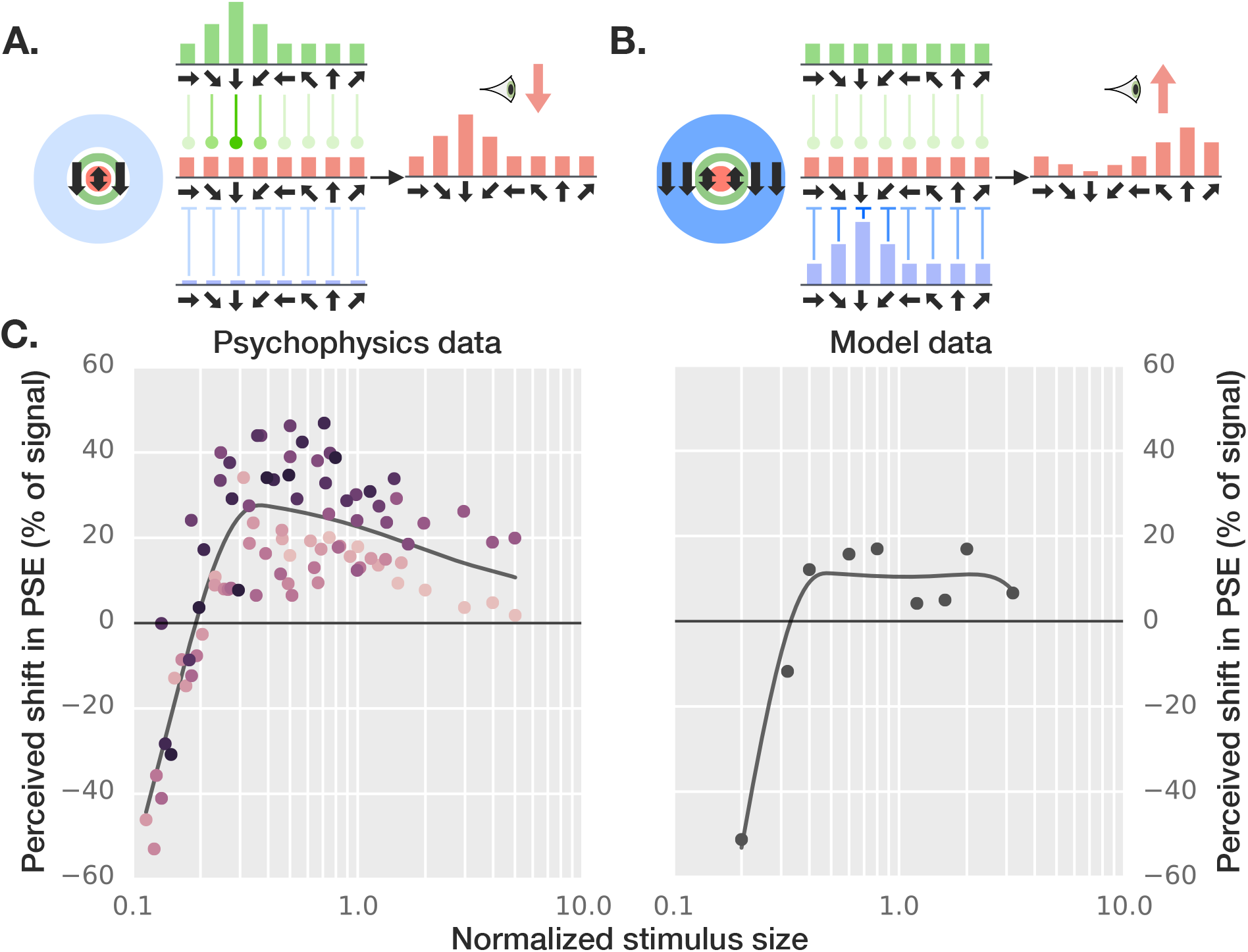
Motion induction. The increasing proportion of the far eCRF activated by larger stimuli explains the shift from assimilation to contrast. **A. Attraction:** When the overall stimulus is small enough, the coherently moving surround dots activate the near eCRF exclusively, leading to motion assimilation (i.e., the center dots’ direction appears the same as that of the surround dots). **B. Repulsion:** Beyond a critical size, activation of the far inhibitory eCRF prevails, which leads to the opposite motion contrast effect (i.e., the center dots’ direction appears opposite to that of the surround dots). **C. Psychophysics vs. model data:** Psychophysics data were digitally extracted from Figure 5 and 6 (Murakami and Shimojo, 1996) and fitted with splines. Both exhibit stronger attraction (negative ordinates) for smaller stimulus sizes, and weaker repulsion (positive ordinates) for larger sizes. Shifts in the point of subjective equality (PSE) were used as a proxy for shifts in perceived motion direction.

We found these results to be consistent with the model (Figure 6C). With a small enough stimulus, the coherently-moving surround dots activate the model near eCRF exclusively (Figure 6A). This leads to a perceptual shift in the direction of the surround, consistent with the analogous case discussed in depth induction. At the population level, facilitation from the near eCRF tends to cause a sharpening in the population response around the surround stimulus value in an otherwise flat population response (as all motion directions are present in the center stimulus). As a result, the center stimulus looks “more like” the surround stimulus. As the stimulus size increases, the coherently-moving surround dots start to activate an increasingly large proportion of the far surround (Figure 6B), which yields the opposite (repulsive) effect. At the population level, suppression from the far eCRF causes a small notch around the surround stimulus value and the center stimulus appears to look “less like” the surround.

### Cooperative activation of the near and far surrounds

Thus far, we have seen that a variety of contextual phenomena can be explained as resulting from a balance between two opposing forces: an attractive force derived from facilitatory mechanisms originating from the near eCRF vs. a repulsive force derived from suppressive mechanisms originating from the far eCRF. This competition can be tipped from attraction to repulsion by increasing the relative contribution of suppressive mechanisms originating from the far eCRF (relative to facilitatory mechanisms from the near eCRF) either by increasing the spatial extent of the stimulus (so as to activate an increasingly large proportion of the far eCRF) or by increasing the similarity between the center and surround stimulus (so as to increase the overlap between center and near surround population responses). However, we reasoned that if a surround stimulus takes on distinct and appropriate values in the near and far eCRFs (which we deem the near and far values), attraction towards the near value could go in the same direction as repulsion from the far value. Thus, the joint activation of the two eCRF sub-regions would cooperate rather than compete, resulting in an even larger perceptual shift compared to what would be achieved by presenting either the near or the far stimulus values alone.

The implication for color perception would be that assimilation, the attraction of the perceived center hue towards a neighboring inducing hue (i.e., the near hue), could be amplified by adding an appropriate outer hue (i.e., the far hue). This idea seems consistent with an “enhanced color shift” illusion discovered by Monnier and Shevell (2003), for which we provide a novel explanation. In the classical color assimilation illusion, a colored test ring (e.g., orange) is presented within a narrow uniform surround (e.g., purple or lime), which then attracts the test ring towards its own hue. This effect was found to be greatly amplified when patterned rings (e.g., alternating, thin rings of purple and lime) at an appropriate spatial frequency and phase were used in place of the uniform colored surround (Figure 1C). Such enhancement has also been documented with achromatic stimuli (Anstis, 2006) and in brightness perception (White, 1979; Anstis, 2006).

Our model provides a simple explanation: As we have established, attraction (i.e., assimilation) towards say purple is caused by the activation of the near surround by a purple stimulus, with respect to a center region coinciding with the test ring. For the appropriate spatial frequency (Figure 7A), the additional lime-colored stimulus activates the far surround, leading to repulsion (i.e., contrast) *away* from lime, thus amplifying the perceptual shift towards purple as purple and lime are roughly perceptual opposites. By reversing the phase of the color grating (Figure 7B), the colors stimulating the near and far eCRFs switch, leading to the same effect in the opposite direction.

**Figure 7.**
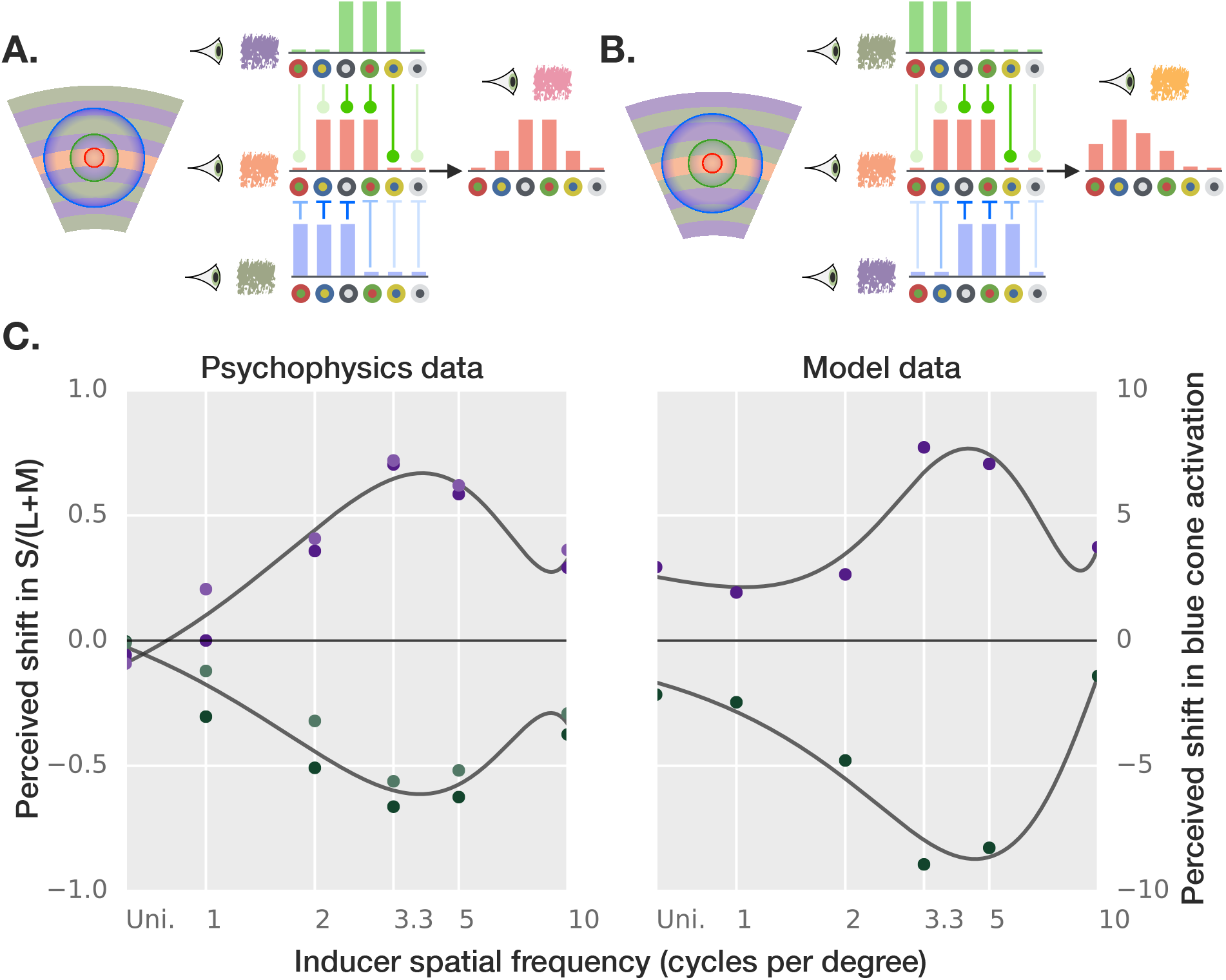
Enhanced color shifts. Cooperative activation of the near and far surrounds explains enhanced perceptual shifts. When distinct and “opposite” hues are used in a patterned surround (or inducer), the resulting shift in color perception of a test hue (here, orange) is amplified relative to a uniform surround of either hue. **A. Shift in one direction:** For the optimal spatial frequency, one surround hue (e.g., purple) overlaps optimally with the near eCRF and the other one (e.g., lime) with the far eCRF. For the right color combination (as here with purple and lime which are “opposite” colors), this results in cooperating perceptual forces: a shift towards purple / away from lime. The colored patches next to the eyes correspond to the color decoded under the ideal observer. **B. Shift in the other direction:** when purple and lime are switched. **C. Psychophysics vs. model data:** Psychophysics data were digitally extracted from Figure 5 (6 minutes test condition) in (Shevell and Monnier, 2005) and fitted with splines. Purple/green dots correspond to condition A/B. ‘Uni.’ stands for a uniform inducer composed of a single hue. For both behavioral and model data, there exists an optimal spatial frequency that maximizes the effect in either direction.

The original psychophysics data (digitally extracted from the “6 min test” curves of Figure 5 from Shevell and Monnier, 2005) and model data are shown in Figure 7C. The model explains the existence of an optimal spatial frequency value, which maximizes the magnitude of the illusion. The spatial frequency of the stimulus controls the strength of the illusion because it determines how cleanly each of the inducing colors (e.g., lime and purple) activate the near and far eCRFs respectively for a CRF centered on the test ring. The model also postdicts that reversing the phases of the color grating leads to an effect with the same amplitude but opposite direction.

Critically, our explanation only depends on the appropriate colors falling within the near and far eCRFs regions; thus, we predict that the periodicity of the inducing stimulus per se is not important, as long as both regions are correctly stimulated. We show this with our own versions of the illusion in Figures S2, suggesting that the illusion is just as strong, if not stronger, when the outer rings are replaced with a single uniform region that activates the far surround optimally (which is not the case for the original stimulus by Monnier and Shevell, 2003).

## Discussion

We have described a computational neuroscience model of recurrent cortical circuits to account for classical (CRF) and extra-classical receptive field (eCRF) effects. The model was constrained by anatomical data and shown in our experiments to be consistent with V1 neurophysiology. In particular, the model unifies several electrophysiology phenomena such as (cross-orientation) normalization within the CRF (Busse et al., 2009) and modulation by the eCRF (including feature-selective suppression, see Trott and Born, 2015) into a computational neuroscience model of contextual integration.

The model further provides computational evidence for the existence of two spatially disjoint eCRF regions with complementary contributions to the CRF (a facilitatory near vs. suppressive far eCRF). In addition, the model makes several novel testable predictions for future electrophysiology experiments. One is the asymmetry between excitation and inhibition in the eCRF: excitation depends on pre-synaptic activity, whereas inhibition depends on both pre-and post-synaptic activities. Another is that short-range connections within a hypercolumn are weakly tuned or untuned, whereas long-range connections across hypercolumns are tuned.

The model distinguishes itself from previous work in succeeding to account for an array of disparate contextual phenomena spanning experimental conditions. Previous computational models have focused on explaining one or a few eCRF phenomena with an emphasis on surround suppression phenomena (see Seriès et al., 2004; Angelucci and Shushruth, 2013; for reviews): Phenomenological models of center-surround processing (Sceniak et al., 2001; Cavanaugh et al., 2002) and other normative models of visual coding (Coen-Cagli et al., 2012; Zhu and Rozell, 2013) have been shown to provide a good fit to single-unit contrast and size tuning responses. Recurrent network models have provided a mechanistic account for some of these phenomena (see Angelucci and Shushruth, 2013; for review) and have even led to testable predictions for single-unit electrophysiology (e.g., Rubin et al., 2015). But, none of these models have been systematically compared to a broad and diverse set of psychophysical experiments.

Furthermore, our model suggests that several contextual phenomena result from not one, but two opposing forces that yield systematic distortions on center population responses: repulsion from the far suppressive eCRF vs. attraction towards the near facilitatory eCRF (see Figure 2B for representative population response dynamics). By revealing commonalities between seemingly disparate perceptual phenomena, the model has helped us establish a novel taxonomy of visual illusions: We have found that the way in which individual stimuli activate these near and far eCRFs (competitively, exclusively or cooperatively; organized by columns in Figure 8) affects the qualitative behavior of the model.

**Figure 8.**
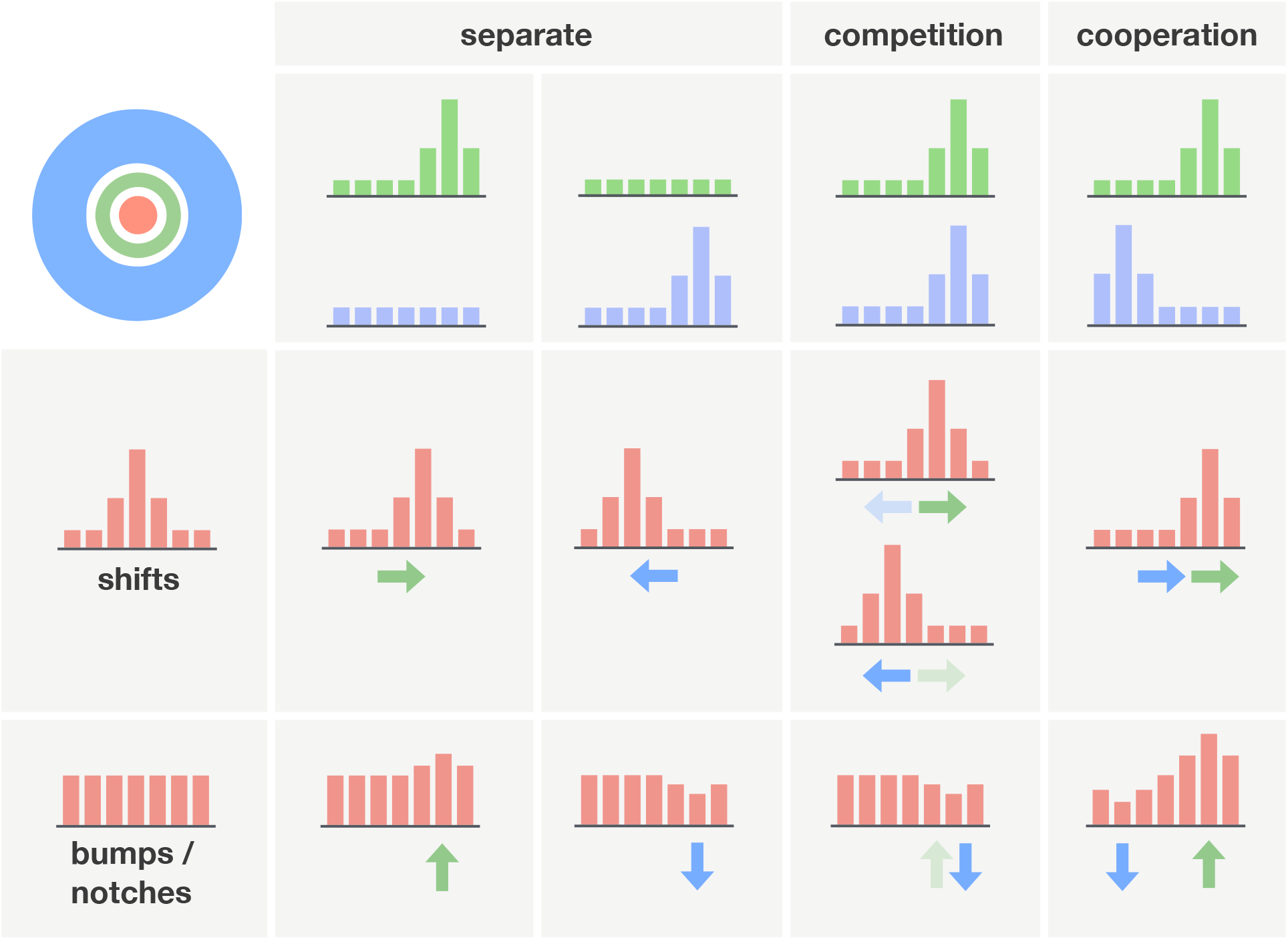
A new taxonomy of contextual phenomena. **Rows:** Contextual phenomena manifest themselves in the model either as (i) shifts with peaked center population response curves (unambiguous stimuli), or (ii) bumps/notches with broad/uniform center population response curves (ambiguous stimuli). **Columns:** Center-surround stimuli activate the near and far eCRFs in three typical ways: (i) either one separately, (ii) both competitively (i.e., near and far eCRFs each induce shifts that tend to stymie each other; green and blue arrows, resp.), and (iii) both cooperatively (i.e., the shifts induced by the near and far eCRFs are in the same direction, amplifying the perceptual shift). See Table S2 for a version of this table populated with representative psychophysics studies for each individual case.

### A novel taxonomy of contextual phenomena

Contextual stimuli that yield competitive activation of the near vs. far eCRFs were found for a set of tilt illusions including orientation (O’Toole and Wenderoth, 1977; Goddard et al., 2008), motion (Kim and Wilson, 1997) and hue (Klauke and Wachtler, 2015; also known as color induction). In these stimuli, the surround spatially overlaps with both the near facilitatory and far suppressive eCRFs – activating them both competitively. Because of the asymmetry between excitation and inhibition in the model, repulsion from the surround stimulus prevails when the center and surround population responses overlap, i.e., when the center and surround stimuli are perceptually similar. Conversely, attraction towards the surround stimulus prevails when such overlap is minimal, such as when the center and surround stimuli are perceptually dissimilar.

Previous authors (Klauke and Wachtler, 2015; Goddard et al., 2008; Kim and Wilson, 1997; Clifford, 2014) have suggested that surround inhibition may be key to explaining the repulsive regime in tilt effects (see Supplementary Discussion for a more in-depth discussion). The proposed mechanisms, which include shifts in neural tuning curves (Klauke and Wachtler, 2015), varying inhibition strength depending on the relative center-surround orientation (Goddard et al., 2008), or recurrent center-surround interactions (Kim and Wilson, 1997) are all consistent with the proposed mechanistic model. In addition, the present study offers a plausible computational explanation for not only the existence of a repulsive regime but also an attractive one for certain classes of stimuli, in agreement with a host of experimental data (O’Toole and Wenderoth, 1977; Goddard et al., 2008; Kim and Wilson, 1997; Westheimer and Levi, 1987).

Another model postdiction is the absence of such attractive regime for contextual stimuli that yield broad-band population responses (arising because of broad neural tuning for the perceptual domain or because the stimulus is inherently ambiguous as in textures with little coherent orientation). For such stimuli, the overlap between center and surround population responses remains large even for maximally dissimilar center and surround stimuli, and the only discernible contextual effect is governed by the repulsive regime. Interestingly, the model achieves its quantitative fit for motion induction experiments via a broadening of neural tuning curves for motion direction compared to orientation, which is consistent with V1 electrophysiology data (Ringach et al., 2002; Albright et al., 1984) (see also Supplementary Discussion and Figure S1 for a more in-depth discussion).

Stimuli that activate exclusively the near or the far eCRF have been used in classical induction experiments in the domain of depth (Westheimer and Levi, 1987) and motion (Murakami and Shimojo, 1996). In the model, consistent with the proposal by Murakami and Shimojo (1993), rescaling a stimulus display (or similarly, varying the relative spacing between center and surround stimuli) yields a reversal from attraction to repulsion. A surround stimulus close to the center or presented at a small scale tends to predominantly activate the facilitatory near eCRF, yielding attraction towards the surround. A surround stimulus farther from the center or presented at a larger scale tends to activate the suppressive far eCRF to a greater extent, yielding repulsion away from the surround.

For the last set of illusions called enhanced color shifts (Shevell and Monnier, 2005), the contextual (or surround) stimulus took on “opposite” optimal values in the near and the far eCRFs. As a result, shifts induced by either region of the eCRF tended to cooperate rather than compete with one another. This resulted in a perceptual shift greater than a purely attractive effect involving only the near eCRF or a purely repulsive effect involving only the far eCRF. In addition, the spatial antagonism of the model eCRF captures the existence of an optimal spatial frequency (and phase) such that a cycle of the surround stimulus coincides maximally with the near and far eCRFs. More generally, the model confirms the consensus that assimilation predominates at higher spatial frequencies and finer scales whereas contrast emerges at lower spatial frequencies and coarser scales (Murakami and Shimojo, 1993; 1996; Monnier and Shevell, 2003; Shevell and Monnier, 2005; White, 1979; 1981; Anstis, 2006).

Shevell and Monnier (2005) have previously modeled enhanced color shifts through an S-cone color opponent model (see Supplementary Discussion). As in our model, such center-surround spatial antagonism results in the existence of an optimal spatial frequency. By design, the model predicts the existence of enhanced perceptual shifts for S-cone stimuli only. However, consistent with our model, we have found such enhanced color shifts to persist for surround stimuli that do not activate S cones (Figure S2).

We have found a further subdivision of the above taxonomy (rows in Figure 8) based on a more detailed characterization of the center stimulus and, in particular, whether it is ambiguous (e.g., incoherently moving random dot or achromatic stimuli) or not (e.g., high-contrast gratings and bars, highly coherent moving random dot or saturated chromatic stimuli). Unambiguous center-surround stimuli yield peaked, narrow population responses (simulation results in Figure 2B-C) across the visual field. The effect of the surround on a peaked center population response is to shift its center of mass, biasing the associated decoded value accordingly (see Supplementary Discussion for a discussion of the evidence of such shifts in neurophysiology studies). The shift is either towards (attraction) or away (repulsion) from the peak of the surround population response depending on whether the net effect of the eCRF is facilitatory (Figure 2B) or suppressive (Figure 2C). Ambiguous center stimuli yield broad-band (or even flat) center population responses (Figure 2D-E). These can be distorted by a peaked surround population in two ways: a bump centered at the surround stimulus value when tuned facilitation from the eCRF prevails (Figure 2D) or a notch at the surround value when tuned suppression does (Figure 2E).

Table S2 shows how the literature fits in the proposed taxonomy. Note that some table entries are missing for certain visual modalities, which suggests more contextual phenomena remain to be found (e.g., cooperative shifts in orientation, which would result in an “enhanced orientation tilt”). Overall, the present study thus provides a vivid example of how computational models may help re-interpret results as well as summarize and integrate disparate phenomena.

### Open questions

Anatomical data to constrain the patterns of recurrent connectivity (both within and across hypercolumns) in the model are scarce. The near and far eCRFs as described in the present model are likely to constitute, at best, coarse approximations corresponding to complex patterns of anatomical connections. In particular, both the spatial extent and the relative strength of the near and far eCRFs relative to that of the CRF were held constant across experiments. Given that the experiments considered throughout spanned a range of visual stimuli across modalities and sizes, it is likely that they recruit neural populations in different cortical areas and visual eccentricities. It is also likely that variations in experimental factors lead to differences in how the center and surround capture attention. We thus expect improvements in the model’s quantitative fit by considering additional parameters to control the spatial extent and the relative strength of the near and far eCRFs (e.g., as done in Goddard et al., 2008).

We also left open on purpose the question of whether the connectivity in the near and far eCRFs would draw on lateral connections within the same area or feedback connections from higher areas (Angelucci et al., 2002a;b; Shushruth and Ichida, 2009). We expect such refinements will be needed to account for some of the electrophysiology phenomena we left aside in the present study including the known contrast dependence of the eCRF size (see Angelucci and Shushruth, 2013; for review) or cross-orientation enhancements (Levitt and Lund, 1997; Sillito et al., 1995).

The neural tuning curves considered in this work (orientation, disparity, motion direction, color opponent) can be found in relatively low-level areas of the visual cortex, such as V1, V2 or MT. Thus, the consistency between model and behavioral data is all the more remarkable as many of the illusions studied here are likely to also involve higher-level visual processes including surface-based and other filling-in processes (Grossberg and Todorović, 1988). Similarly, considering higher-level tuning curves such as those found for V4/PIT hue-selective neurons (as opposed to color-opponent V1 neurons considered here, see Conway et al., 2007) would also improve the fit with experimental data (though hue tuning remains controversial, see Mollon, 2009; Conway, 2009).

Similarly, we have assumed for simplicity that the near eCRF is circular (i.e., isotropic with respect to the topography of the visual field). There is, however, evidence for anisotropies in the pattern of horizontal connections between cortical columns (as orientation-tuned cells tend to be more often connected when they share the same selectivity and their CRFs are aligned along an axis parallel to their preferred orientation, see Bosking et al., 1997). There is also more direct evidence for anisotropies in the shape of the eCRF (i.e., various elongations over a wide range of orientations and widths, see Tanaka and Ohzawa, 2009). The function of these anisotropies has been attributed to the computation of higher-order features (including contrast-or texture-defined boundaries) as well as contour integration and pop-out (Stemmler et al., 1995; Hess et al., 2003; Tanaka and Ohzawa, 2009). Future work should test whether these phenomena can be accounted for with a model extension that incorporates such eCRF anisotropies.

We speculate that the computational mechanisms revealed by the contextual illusions studies here play a key role in shaping invariant population codes for object constancy. We have obtained preliminary results suggesting that tuned suppression from the far eCRF may improve the accurate decoding of surface reflectances across changes in illumination (i.e., color constancy; see Mély & Serre, abstract presented at the 2015 Vision Science Society meeting), by helping to discount undesirable variations in center population responses caused by changes in the light source. (This is reminiscent of a color constancy algorithm by Land and McCann (1971) known as the Retinex.) This raises the intriguing possibility that at least some of the mechanisms unraveled here may support other forms of perceptual constancy beyond color. Further work will be needed to quantify how object transformations such as changes in illumination or depth affect neural population responses tuned to orientation or binocular disparity and what computational mechanisms are needed to help discount these nuisances. Nonetheless, the ability of the model to account for the variety and complexity of contextual illusions provides computational evidence for a novel canonical cortical circuit shared across visual modalities.

## Materials and Methods

Additional methods may be found in Supplemental Materials and Methods.

### Model connectivity

A column centered at location (*x,y*) contains a complete set of N units with CRFs centered at (*x,y*) and tuning values covering the full range *θ_k=1…N_* (e.g., orientation tuning curves are regularly centered at values *θ_k_*∈[0,180°]). Tuning curves are idealized – either bell-shaped for disparity (Cumming and Parker, 1997), motion direction (Albright et al., 1984) and orientation (Ringach et al., 1997) or monotonic for color opponency (Johnson et al., 2001).

Each unit (*x,y,k*) receives excitation 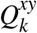, assumed to be weakly tuned and originating from within the same hypercolumn:

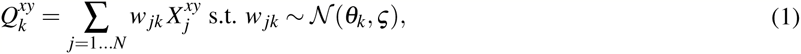

where *w_jk_* corresponds to excitatory weights between units *k* and *j* (with selectivity *θ_k_* and *θ_j_*, respectively) and 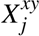 to input activity at location (*x,y,j*). We assume these weights to be normally distributed, centered at a target unit tuning preference *θ_k_* with standard deviation *ç*. Some tuning (albeit weak) is necessary in order to prevent intra-columnar excitation from flattening the population responses to well-defined stimuli. In the color domain, we consider color-opponent model units with monotonic tuning curves. Instead of drawing weights from a normal distribution, which only makes sense for bell-shaped tuning curves, we set 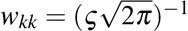 and *w_jk_* = const. (when *j ≠ k*; under the constraint that the weights sum up to 1).

Each unit (*x,y,k*) also receives some inhibition *U^xy^*, assumed to be untuned and originating from within the same hypercolumn:

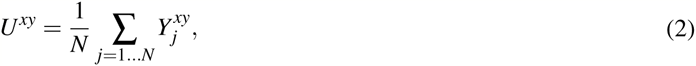

where 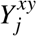 is the output activity of unit *j* at location (*x,y*). Because the local inhibition is untuned, its strength is independent of a unit selectivity *θ_k_*, and we drop the subscript k for simplicity.

Furthermore, unit (*x,y,k*) also receives tuned excitation 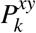 from other units with the same selectivity *θ_k_* that are located within its near eCRF *ℕ^xy^*, defined relatively to position *(x,y)*:

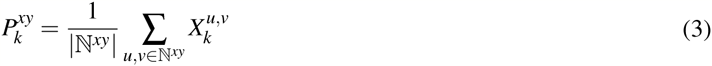

Similarly, unit (*x,y,k*) also receives tuned inhibition from 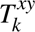 other units with the same selectivity *θ_k_* that are located within its far eCRF *Ϝ^xy^*, defined relatively to position (*x,y*):

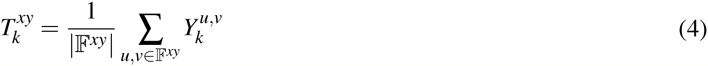

### Neural field model

Neural field dynamics obey the following equations:

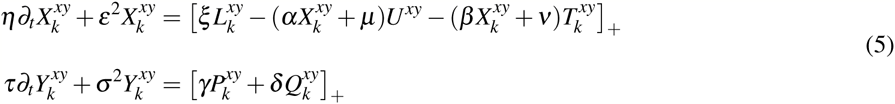

where the feed-forward input 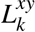 drives every unit (*x,y,k*) across the visual field; each is represented by its recurrent input 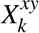 and output 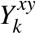 The steady-state solution (analytic form given in Supplementary Information, Equation S1) is computed using numerical integration (with convergence typically taking ~ 50 iterations). Population responses at the steady-state 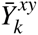 are a very nonlinear function of the model input 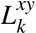.

For each unit, the steady-state input and output are given by 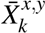 and 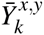 resp. Due to the rectifying non-linearity in the dynamics (Equation), at steady-state, 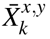 and 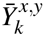 can either be equal to zero, or to the values below:

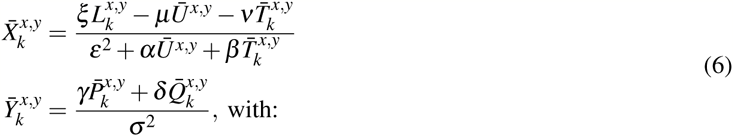

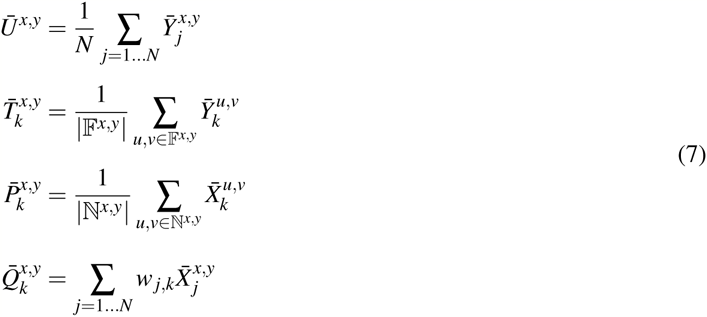

### Tuning curves

We considered two kinds of tuning curves: bell-shaped (orientation, motion direction, binocular disparity) and monotonic, non-saturating tuning curves (color). All tuning curves were normalized, i.e., the maximum unit activity was set to be equal to 1. For non-angular variables (e.g., disparity), bell-shaped tuning curves were parametrized as Gaussian functions:

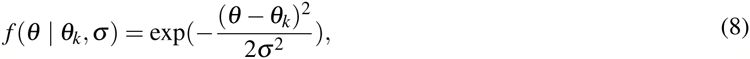

with preferred stimulus value *θ_k_* and tuning bandwidth σ. When the variable was circular (e.g., orientation, motion direction), we modeled the tuning curve as a von Mises function instead:

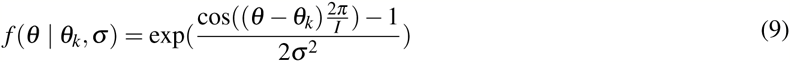

where *I* indicates the length of domain of the tuning curve (e.g., π for orientation vs. 2π for direction). We generally sampled on the order of 30 tuning curve centers regularly spaced in the domain of the considered visual modality. We found that the number of tuning curve centers considered did not impact our results as long as it was large enough.

Monotonic, non-saturating tuning curves for color were derived by converting stimuli to idealized cone responses first, which were then mapped to opponent color channels similarly to Zhang et al. (2012). These included red-on/green-off (*R^+^G^−^*), green-on/red-off (*G^+^R^−^*), blue-on/yellow-off (*B^+^Y^−^*), and yellow-on/blue-off (*Y^+^B^−^*), alongside with a pair of luminance-sensitive channels, selective for lighter (*Wh^+^Bl^−^*) and darker (*Bl^+^Wh^−^*) stimuli.

### Model parameters

All circuit parameters were held constant in all comparisons with psychophysics data. They were determined a priori in order to reproduce key neurophysiology data (see Supplementary Experiments) and were held constant for all visual modalities except color because of a qualitative difference in tuning curve (see Equation 1). In all subsequent experiments, only two variables were allowed to vary: the stimulus scale and the tuning bandwidth for model units.

The stimuli used in psychophysics studies varied greatly – recruiting neural populations subtending a wide range of CRF (and eCRF) sizes and eccentricities, possibly spanning different visual areas. Rather than adjusting the size of the model CRFs and eCRFs for individual experiments, which would have required structural changes to the model, we instead varied the stimulus scale. Because the connectivity between model hypercolumns was held fixed, this is somewhat akin to varying the magnification factor in the model. Critically, this yielded broad estimates for CRF (and eCRF) sizes within a biologically realistic range (from a fraction of a degree of visual angle to a couple of degrees). The width of the idealized tuning curves which is common to all model units was optimized separately for each experiment (see Supplementary Materials and Methods for details). We have confirmed that our key model predictions were robust over a range of these parameter values.

### Ideal neural observer model

We used an ideal neural observer model to map model population responses to decoded sensory variables, which can then be compared to behavioral judgements collected experimentally. We used a population vector model (Georgopoulos et al., 1986), in which each unit votes for its preferred sensory value in proportion to its activity (normalized by the summed activities of all units within the same column). This model is not appropriate for color because of the tuning along opponent color pairs rather than a hue angle. Instead, we used cross-validated ridge regression to decode the sine and cosine of hue.

## Acknowledgments

We would like to thank Dr. Drew Linsley for helpful comments. This work was supported by DARPA young faculty award [grant number YFA N66001-14-1-4037] and NSF early career award [grant number IIS-1252951] to TS.

